# Context-dependent structure formation of hairpin motifs in bacteriophage MS2 genomic RNA

**DOI:** 10.1101/2024.04.17.589867

**Authors:** Veronika Bukina, Anže Božič

## Abstract

Many functions of ribonucleic acid (RNA) rely on its ability to assume specific sequence-structure motifs. Packaging signals found in certain RNA viruses are one such prominent example of functional RNA motifs. These signals are short hairpin loops that interact with coat proteins and drive viral self-assembly. As they are found in different positions along the much longer genomic RNA, the formation of their correct structure occurs as a part of a larger context. Any changes to this context can consequently lead to changes in the structure of the motifs themselves. In fact, previous studies have shown that structure and function of RNA motifs can be highly context-sensitive to the flanking sequence surrounding them. However, in what ways different flanking sequences influence the structure of an RNA motif they surround has yet to be studied in detail. We focus on a hairpin-rich region of the RNA genome of bacteriophage MS2—a well-studied RNA virus with a wide potential for use in biotechnology—and systematically examine context-dependent structural stability of 14 previously identified hairpin motifs, which include putative and confirmed packaging signals. Combining secondary and tertiary RNA structure prediction of the hairpin motifs placed in different contexts, ranging from the native genomic sequence to random RNA sequences and unstructured poly-U sequences, we determine different measures of motif structural stability. In this way, we show that while some motif structures can be stable in any context, others require specific context provided by the genome. Our results demonstrate the importance of context in RNA structure formation and how changes in the flanking sequence of an RNA motif sometimes lead to drastic changes in its structure. Structural stability of a motif in different contexts could provide additional insights into its functionality as well as assist in determining whether it remains functional when intentionally placed in other contexts.

**STATEMENT OF SIGNIFICANCE:** RNA motifs are groups of related RNAs that possess similar sequence and/or structure and consequently assume similar functions. Despite their similarities, these motifs are often only a small part of larger RNA molecules, situated in various contexts provided by the surrounding (flanking) sequences. How the nature of the flanking sequence influences the structure of a motif it surrounds is a fundamental yet underexplored question. We systematically study context dependence of several *hairpin motifs* in the genomic RNA of bacteriophage MS2 which act as packaging signals, indispensable for virus assembly. We show that while some motifs fold into the correct structure no matter the nature of their context, others require the specific context provided by the genomic RNA.

## INTRODUCTION

Ribonucleic acid (RNA) takes on myriad functions in the biological environment: from a messenger RNA (mRNA) between the genome and the protein product to its various non-coding roles as transfer RNA (tRNA), ribosomal RNA (rRNA), microRNA (miRNA), and long non-coding RNA (lncRNA), to mention just a few [1, 2]. Importantly, for a large number of bacterial, plant, animal, and human viruses alike, single-stranded RNA (ssRNA) assumes the role of their genomes [3], a unique feature among living organisms. Many functions of RNA are thus carried out not only at the level of its primary *sequence* of nucleotides (A, C, G, and U) but also at the level of its secondary and tertiary *structure* [4–6], whose formation is a consequence of base-pairing and tertiary interactions between the nucleotides [7–9]. This is no less true for ssRNA viruses, where the requirement for the genomic RNA (gRNA) to encode both the protein product as well as functionally important local and global structural elements leads to a high density of functions encoded in viral gRNA [10–15].

Groups of related RNAs that possess similar sequence and/or structure are termed *RNA motifs* [16, 17] and often carry out similar functions. Of these, the relatively simple *hairpin motif* stands out as it can guide RNA folding, protect mRNA from degradation, act as a substrate for enzymatic reactions, or serve as a recognition motif for RNA binding proteins (RBPs) [18]. It is in this latter function that hairpin motifs can often be found in many positive-strand ssRNA (+ssRNA) viruses, where they function as *packaging signals* (PSs). The PS motifs are high-affinity binding sites for the viral capsid (or nucleocapsid) proteins that distinguish viral gRNA from cellular RNAs and drive the virion assembly process [19, 20]. These motifs have been proposed to play a role in different stages of life cycles of some viruses [15], and the emergence of a robust RNA hairpin packaging cassette analogous to viral PSs has also been observed in the course of a laboratory evolution of a bacterial enzyme nucleocapsid packaging its own mRNA [21].

Many simple +ssRNA viruses consisting solely of a proteinaceous capsid and viral gRNA have been shown to utilize one or more PS motifs to selectively package their genomes [22–28], even though the presence and functionality of these motifs are often difficult to establish [20]. One of the most well-studied viral systems in this respect is bacteriophage MS2, a small +ssRNA virus that infects bacteria [26, 27, 29–35]. Based on theoretical analyses, its self-assembly has been proposed to follow a Hamiltonian path [28, 36] which hinges on the presence of numerous PS motifs along its genome [37, 38]. Indeed, different experimental studies found numerous *putative* PS sites in the MS2 gRNA [31, 32]. Moreover, asymmetric cryo-EM reconstruction of the MS2 genome inside the capsid has shown that the RNA forms a branched network of hairpin motifs, almost all of which are located near the inner surface of the capsid [33]. Conserved motifs of RNA–coat protein interactions in these studies indicate that any restrictions on the phage MS2 PS motifs are mostly on their structure, with only a few unpaired nucleotides in the structure of the hairpin motif being constrained [30–32].

Functional structure of a PS motif always has to form in the presence of a *flanking RNA sequence* provided by the surrounding gRNA. In general, flanking sequences— RNA sequences either up- or downstream of a sequence or structure of interest—have been shown to have a profound effect on the functionality of the region they surround in a wide array of RNAs. These range from miRNA processing [39, 40] and binding and regulation of RBPs [41, 42] to various functional parts of the RNA genomes of viruses, such as the frameshifting element (FSE) in the genome of severe acute respiratory syndrome coronavirus 2 (SARSCoV-2) [43], the transcription consensus sequence of mouse hepatitis virus [44], the hepatitis delta virus ribozyme [45], and a sequence-nonspecific hairpin that enhances replication of turnip crinkle virus satellite RNA [46]. Importantly, in some cases, flanking *sequences* have been shown to play an important role in the formation of RNA *structures*. For instance, CAG-repeat regions in huntingtin mRNA were shown to adopt distinct structures depending on the flanking sequence context [47], while the FSE found in SARS-CoV-2 gRNA has been demonstrated to require native context sequence of the genome to fold into the correct structure [43]. Moreover, flanking sequences have been shown to stabilize the transcript and aid proper folding and dynamics of an RNA thermometer [48] and influence the cotranscriptional folding kinetics of a 99-nucleotide (nt)-long region in the hepatitis delta virus ribozyme [45]. Changes in context are also an important consideration in the design of synthetic RNA motifs as they can lead to unexpected divergences between the designed and the actual function of the motifs [49].

While different studies have shown that RNA folding and function are highly context-sensitive, few of them have systematically explored in detail how the *nature* of the flanking sequence and *changes* in it influence the structure of an RNA motif they surround, and we set out to tackle this challenge in our work. To achieve this, we focus on the influence of different flanking sequences on the structure formation of 14 hairpin motifs in an ∼800-nt-long region of the MS2 bacteriophage gRNA. MS2 gRNA is particularly well-suited for our study, as more than 50 hairpin motifs have been identified to bind to virus coat proteins [31], but only a few of them have been firmly established to play the role of a PS motif [30–32]. We define the native (target) structures of the 14 motifs based on the experimental study of Rolfsson *et al*. [31] and use a combination of computational secondary and tertiary structure prediction together with several structural stability scores to study how not only the presence of a flanking sequence but also its nucleotide composition influence the structure of hairpin motifs. Our results show that while some motifs will always form into the native structure regardless of the presence or absence of a flanking sequence, others require the specific context provided by the surrounding RNA genome to be able to fold into the correct structure. Implications of our study reach beyond the stability of hairpin motifs and PS structural elements in MS2 and related +ssRNA bacteriophages [50–53]: stable local structural elements positioned in a larger RNA context sequence are also required for RBP binding [54], correct cleavage of mitochondrial RNA at tRNA sites [55–57], in the design of modular and reusable RNA regulatory devices [58], and in bioengineered protein-RNA complexes where phage PSs are inserted into a non-native RNA context [51, 59–61].

## METHODS

### MS2 gRNA hairpin motifs

Apart from the well-known TR PS motif in MS2 gRNA, we consider 13 other hairpin motifs in its vicinity which have been defined with varying degrees of certainty [30–32]. These motifs are located in the region of the MS2 genome between nucleotide 1457 and nucleotide 2171 and span anywhere from 11 nt to 22 nt in length. This region of the MS2 genome spans the end of the coat protein coding region, start of the replicase protein coding region, and the entirety of the lysis protein coding region which overlaps with the other two regions [34].

The 14 hairpin motifs that we study are characterized by a central hairpin loop with a size of 3–8 nt and a surrounding stem. Few motifs have an additional single-nucleotide bulge in the stem. Figure S1 in the Supplementary Material highlights these motifs in the relevant region of the MS2 gRNA, and their sequences and secondary structures are listed in Table S1 in the Supplementary Material. The notation we use for motif labelling follows the work by Rolfsson *et al*. [31].

We note that some of the 14 motifs have also been identified in a separate cryoEM-based study by Dai *et al*. [32], in particular the motifs SL-3, SL-1, TR, SL+1, and SL+4. While the size of these motifs varies slightly from those listed in Table S1, their predicted secondary structures match the ones proposed by Rolfsson *et al*. [31]. Exception is the SL+1 motif, for which Dai *et al*. [32] predict a structure with a single-nucleotide bulge similar to the structure of the TR motif. Keeping this in mind, we will refer to the sequence-structure hairpin motifs presented in Table S1 as “native” or “target” motifs, in order to distinguish them from the motif structures which will arise during their folding in the presence of different flanking sequences.

### Flanking sequences

To study how the structures of the hairpin motifs change as their flanking sequence—sequence immediately up- and downstream of the motif sequence—is modified, we surround the motif sequence of length *N* by different flanking sequences of length *L* on each side of the motif, yielding an RNA sequence of total length *N* + 2*L* (Fig. 1). The flanking sequence itself is either *(i)* generated randomly (with each nucleotide A, C, G, and U having equal probability); *(ii)* taken from the corresponding genomic sequence of MS2 (GenBank: V00642.1); or *(iii)* composed solely of U nucleotides (poly-U flanking sequence) so that it does not form a secondary structure on its own. When considering random flanking sequences, we generate 500 different flanking sequences on each side of the motif.

**Figure 1.**
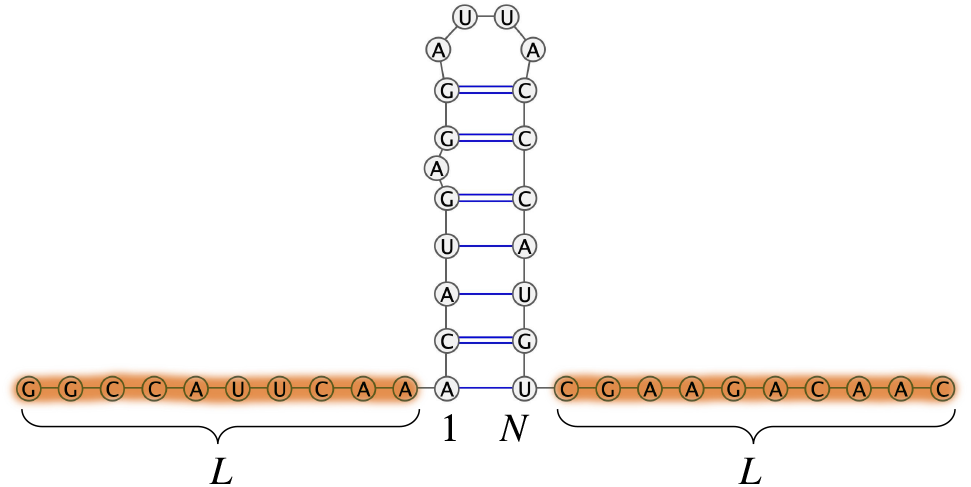
Sketch of flanking sequence insertion up- and downstream of a hairpin motif of length *N*. Two flanking sequences of length *L* (highlighted) are inserted to the left and right of a chosen motif. The joint sequence of total length *N* + 2*L* is then used to determine its secondary and tertiary structure, based on which we analyze the structure of the motif region.

### Secondary structure prediction

To predict the secondary (2D) structure of a motif and its flanking sequence, we use ViennaRNA software (v2.5.1) [62]. We use the standard settings with an additional flag --noLP, which forbids structures with lonely pairs (helices of length 1). Secondary structures are determined at temperature *T* = 37 ^*°*^C and monovalent salt concentration *c*_0_ = 1 M.

For each sequence, we obtain the pair probability matrix *P*_*ij*_, which gives the probability that nucleotides *i* and *j* form a base pair. The probability that a nucleotide is unpaired is stored in the diagonal elements, i.e., 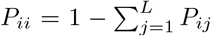, which is important for the calculation of different structural stability measures. Since the energy landscape of RNA structures is typically quite rugged and shallow, the minimum free energy state is not necessarily the biologically relevant state [63, 64]. We thus also generate 3 *×* 10^3^ secondary structures sampled from the thermal ensemble for each sequence and use them in our analysis.

### Tertiary structure prediction

To predict the three-dimensional (3D) structure of a motif and its flanking sequence, we use the sequence-dependent version of oxRNA software [65–67]. We use virtual move Monte Carlo (VMMC) algorithm implemented in the software to explore the configuration space of RNA structures and let the simulation run for 2 *×* 10^7^ steps. After equilibrating the system for 10^5^ steps, we use the main part of the simulation run to observe transitions between different RNA structures. We run 40 replicas for each studied case, with starting configurations randomly sampled from among the structures obtained during a single run at a high temperature (*T* = 50 ^*°*^C). The temperature and salt concentration of the simulations are set to match those used in ViennaRNA (*T* = 37 ^*°*^C and *c*_0_ = 1 M, respectively). Due to the large number of motifs and flanking sequences studied, we limit ourselves to flanking sequences of length *L* = 30 nt, as we observe from the secondary structure prediction that this length suffices to induce structural changes and is at the same time still computationally feasible.

#### Extracting secondary structures

We use the accompanying tools in oxRNA to monitor the energies of the 3D structures as well as the number and nature of the bonds between nucleotides. We sample the 3D structures every 5000 steps, resulting in 4000 structures from each VMMC run. We use the information on the strength of the hydrogen bonds between nucleotides to define whether they form a base pair or not, and in this way determine the secondary structure representation of the 3D RNA molecule. The cut-off for the formation of a base pair in oxRNA is set as the energy of the hydrogen bond between nucleotides *i* and *j* being 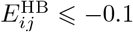 in simulation units (corresponding to 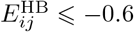 kcal*/*mol) [66, 68].

### Measures of motif structural stability

A sequence-structure motif ℳ (*m, N*) is a part of a larger RNA sequence of length *K* that starts at position *m* and spans *N* nucleotides. We require that the native (target) secondary structure of the motif is complete, i.e., that no part of the motif sequence forms base pairs outside of it. To analyze how the addition of different flanking sequences around a motif changes its structure, we adopt several different measures that operate on its secondary structure representation [69, 70]: *(i)* structure probability 𝒫 (ℳ), *(ii)* ensemble defect ED(ℳ), and *(iii)* Shannon entropy SE(ℳ).

#### Structure probability

A simple yet powerful measure of motif stability is the likelihood of its *exact* structure occurring at equilibrium, 𝒫 (ℳ) [70]. The value of structure probability is obtained from the thermal ensemble of RNA structures, counting the number of occurrences of the exact (target) motif and dividing it with the number of all structures in the ensemble. As already mentioned previously, we typically consider for each case a sample of 3 *×*10^3^ secondary structures from the thermal ensemble, or a sample 4 *×* 10^3^ 3D structures from each of the 40 replicas of VMMC runs.

#### Ensemble defect

Another measure for the stability of a motif is the ensemble defect (ED), introduced by Dirks *et al*. [71], which gives the weighted average number of incorrect base pairs in the ensemble given a target structure. An ED of 0 represents an ensemble which always acquires the target structure, and an increase in ED indicates an increased proportion of alternative folds in the ensemble. Since we are interested in a particular motif ℳas a part of a longer sequence—whose structure is not of interest to us—we need to restrict the definition of ED to the motif only:

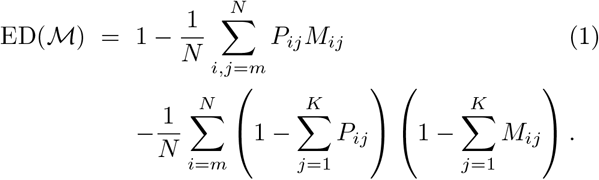

Here, we have introduced the base pair matrix of the motif as *M*_*ij*_, whose elements are *M*_*ij*_ = 1 if the bases *i* and *j* form a base pair in the motif and are equal to zero otherwise; the diagonal elements *M*_*ii*_ contain the information on unpaired nucleotides. The definition in Eq. (1) can be recast in terms of the suboptimal structures sampled from the thermal ensemble, which is the version we use to determine ED on the secondary structure representation of 3D structures since the base pair matrix is not available in that case.

#### Shannon entropy

Lastly, we make use of Shannon entropy (SE), which is a measure of how reliably a given base pair is formed in a thermal ensemble of RNA structures. A low SE means that an RNA sequence reliably forms into a given structure with few deviations, while a high SE implies that the RNA samples many different structures and base pair patterns. Since we are again interested only in the stability of the motif region, we limit our definition of SE to the motif:

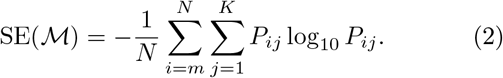

We note that both log_2_ and log_10_ are used in the literature in the definition of Shannon entropy; we opt for the logarithm with base 10 as it seems to be the one used more often in recent works [72–74]. We determine Shannon entropy only for predictions based on secondary structure, as only these allow us to obtain the pair probability matrix.

### Gaussian mixture models for distributions of structural measures of random flanking sequences

When a motif is surrounded by a random flanking sequence, we observe in the structural measures that the presence of the native motif structure is gradually reduced at the expense of non-native structures, with the extent of the change depending on the length and composition of the flanking sequence. In order to separate, at least approximately, the contributions of different populations of motif structures to structural measures, we apply a Gaussian mixture model [75] to the distributions of these measures across a sample of 500 random flanking sequences. Typically, mixtures of 2 or 3 components suffice to describe the ensemble, as determined by the minimum of the BIC curve [76]. In order to keep comparison simple, we limit ourselves to at most 3 components and assign to each an average of a given quantity over the sequences within the component; we also retain the corresponding weight (proportion) of the component. Figure S2 in the Supplementary Material shows a detailed example of this approach.

### Shuffling motif sequences

As a point of additional comparison between different motifs and their structures, we compare the folding free energies of the thermal ensemble of structures stemming from either the motif sequence or its mono- and dinucleotide shuffled versions. We use a *k*-let preserving shuffling algorithm implemented by uShuffle [77] to shuffle the original motif sequences while exactly preserving 1-let (mononucleotide) and 2-let (dinucleotide) frequencies.

## RESULTS

The MS2 hairpin motifs are an integral part of the larger sequence and structure of the genome, even though they represent defined sequence-structure motifs in and of themselves. As a first step in our analysis, we determine several measures of structural stability—structure probability 𝒫, ensemble defect ED, and Shannon entropy SE (see Methods)—for all 14 hairpin motifs (Fig. S1 and Table S1) without any modifications to their flanking sequence. We do this both by predicting the secondary structure of solely the motif sequence on its own, comparing the resulting structure to the native (target) one, and by folding the motif sequence together with the entirety of the MS2 gRNA, again comparing only the structure of the PS motif to the predicted one.

The different structural measures for these two cases are shown in Fig. 2, where we can already observe that there are significant differences in how reliably the 14 motif sequences fold into the correct structure when they are folded alone compared to when they are a part of the entire genome. For most motifs, the presence of MS2 gRNA— their native context—improves all structural measures (shaded regions in panels (a)–(c) of Fig. 2): Firstly, they exhibit an increase in structure probability 𝒫 (panel (a)), meaning that the *exact* native (target) structure occurs more likely in the thermal ensemble when the motif is a part of the MS2 gRNA. They also show a decrease in the ensemble defect ED (panel (b)) in the presence of gRNA, which implies that apart from the exact structure being more likely, other structures that the motif sequence assumes in the thermal ensemble are close to the native one. Lastly, the Shannon entropy SE (panel (c)) of the folded motifs also decreases in the presence of gRNA, which means that fluctuations around the native structure are small.

**Figure 2.**
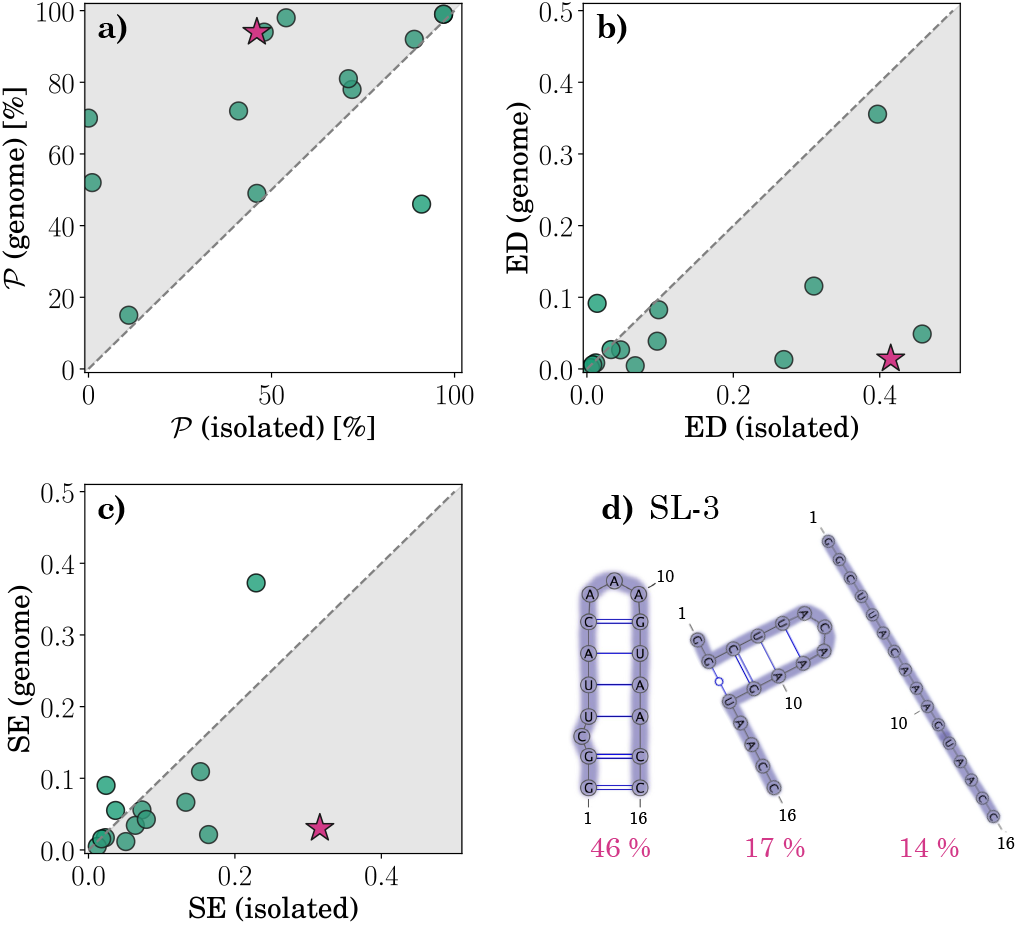
**(a)** Structure probability *P*, **(b)** ensemble defect ED, and **(c)** Shannon entropy SE for the secondary structures of the 14 MS2 hairpin motifs considered in this work. Shown are the values for isolated motifs (i.e., motif sequence folded by itself) and the motifs folded in the presence of the entire MS2 gRNA. Shaded areas show the regions where the structural stability measures are improved in the presence of gRNA. Magenta star denotes the hairpin motif SL-3 (Table S1). **(d)** The three most commonly occurring structures of the motif SL-3 when folded on its own, together with the percentages of their occurrence. When folded in the presence of the entire MS2 genome, only the leftmost—most common—structure is present.

In several cases, the presence of MS2 gRNA increases the probability of the occurrence of the native motif structure in the thermal ensemble by several fold. For instance, the sequence of the motif SL-3 folds on its own into the native (target) structure only in 𝒫= 46% of the structures, while the other two most commonly occurring structures (representing 17% and 14% of the cases, respectively) are far from the native one (Fig. 2d). However, when the motif sequence is folded in the context of the entire MS2 gRNA, the hairpin motif assumes the native structure in 𝒫 = 94% of the cases (cf. Table S2 in the Supplementary Material). Only in a single case of the motif SL+3 does the presence of gRNA actually reduce the likelihood of the native motif structure occurring (Table S2).

### Context provided by MS2 gRNA can improve structural stability of hairpin motifs

In some very clear cases, which include motifs SL-3, IX, and XIV (and partially also III), the presence of the flanking sequence provided by the MS2 gRNA is required in order for the motif itself to form the correct structure; an example of the motif SL-3 is shown in Fig. 3. We can observe that when the motif sequence is either folded alone or when it is surrounded by a short (*L* = 30 nt) poly-U sequence (which does not form secondary structure on its own), the native (target) motif structure is present in less than 50% of the cases (cf. also Fig. 2d). This is the same when the motif is surrounded by a short random flanking sequence: only a very small proportion of the 500 random flanking sequences considered leads to the correct folding of the SL-3 motif. However, when the short flanking sequence is taken from the MS2 gRNA or when the motif is folded in the presence of the entire genome, the probability of the correct motif structure occurring rises to almost 100%, based on the prediction of its secondary structure. The same trend is observed for the predictions based on tertiary structure. Moreover, the ensemble defect of the motifs in the presence of non-genomic flanking sequences is fairly high, indicating that the structures that the motif sequence folds into are distant from the native one. Similarly, their Shannon entropy is also high, meaning that the base-pairing pattern in the misfolded structures varies to a large extent. On the contrary, in the presence of the gRNA flanking sequences, both ED and SE are low, indicating a stable motif structure corresponding to the native (target) one.

**Figure 3.**
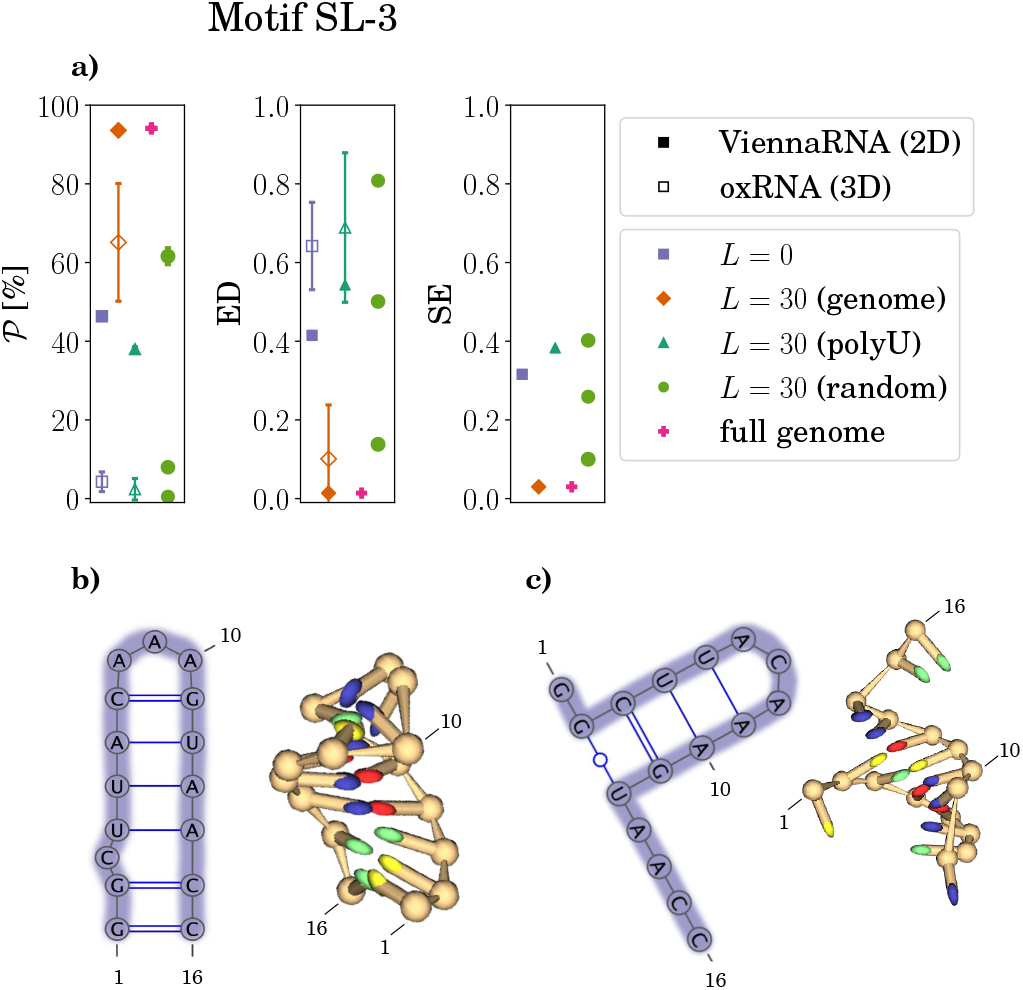
**(a)** Structure probability, ensemble defect, and Shannon entropy of the SL-3 hairpin motif structure as determined by ViennaRNA (full symbols) and oxRNA (empty symbols). The values are shown for the motif in absence of flanking sequence (*L* = 0) as well as in the presence of different flanking sequences—genomic, poly-U, and random—of length *L* = 30 nt. Secondary structure prediction also includes the motif in the context of the full MS2 gRNA. Size of the symbols for random flanking sequences corresponds to the weight of the corresponding component of the Gaussian mixture model (see Methods and Fig. S2 in the Supplementary Material). **(b)** Secondary and tertiary structure of the native (target) SL-3 hairpin motif. **(c)** Secondary and tertiary structure of the most common misfolded state of the SL-3 sequence.

Similar observations can be made for two more hair-pin motifs, IX and XIV (Fig. S3 in the Supplementary Material). In both of these cases, the presence of the MS2 gRNA flanking sequence is required for the native motif structure to form. While these trends are observed in all three motifs based on the predictions of secondary structures obtained either directly from ViennaRNA or extracted from the 3D structure predicted by oxRNA, the exact values of the different structural stability measures can differ between the 2D and 3D predictions. We note that this is often (but not always) a consequence of the secondary structure being an inexact and not easy to extract representation of the full 3D structure; we comment further on this observation in the Discussion.

### Some motif sequences always lead to stable hairpin structures

In contrast to previous examples, structures of some hairpin motifs—notably, SL-1, TR, IV, and VIII (and partially II)—tend to correspond to the native (target) ones regardless of the context they are folded in. An illustrative example is the motif SL-1, for which Fig. 4 shows how the probability of the motif occurrence, its ensemble defect, and Shannon entropy are affected when different flanking sequences are present. We can observe that the motif sequence folded on its own or in the presence of either poly-U or genomic flanking sequence assumes the native (target) structure with high probability, 𝒫 ≳75%. Random flanking sequences can still lead to a lower probability of motif occurrence and an increase in both ensemble defect and Shannon entropy (Table S2): For 30% of the 500 random flanking sequences, the motif sequence folds into the native structure, while for the other 70%, the motif sequence typically folds into different structures.The ED and SE values of the latter, however, show that these structures are still fairly close to the native one.

**Figure 4.**
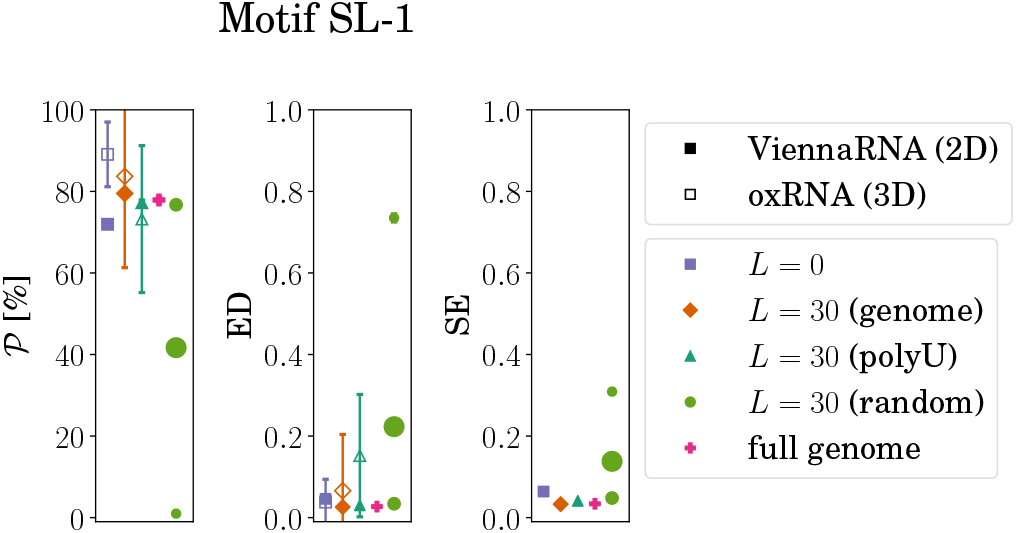
Structure probability, ensemble defect, and Shannon entropy of the SL-1 hairpin motif structure as determined by ViennaRNA (full symbols) and oxRNA (empty symbols). The values are shown for the motif in absence of flanking sequence (*L* = 0) as well as in the presence of different flanking sequences—genomic, poly-U, and random—of length *L* = 30 nt. Secondary structure prediction also includes the motif in the context of the full MS2 gRNA. Size of the symbols for random flanking sequences corresponds to the weight of the corresponding component of the Gaussian mixture model (see Methods and Fig. S2 in the Supplementary Material).

The TR motif (Fig. S3) is a rare—although given its importance, perhaps not unsurprising—case where all flanking sequences, including the *majority* of the 500 random flanking sequences, lead to a reliable formation of its native secondary structure. On the other hand, in other hairpin motifs that we study, adding a random flanking sequence is usually likely to reduce the chance of correct folding to at least some extent (Table S2).

### Determination of motif stability depends on the accuracy of target structure

Sometimes we also observe that the motif structure is found to be stable when folded on its own or in the context of poly-U sequence but is, on the other hand, destabilized when put in the context of the native MS2 gRNA—such is the case for motifs SL+3 and XIII (Fig. S3). In other cases, the native (target) motif structure does not occur to any large extent (as determined by the three measures of structural stability), no matter what context it is in (motifs II, III, and SL+1; cf. again Fig. S3).

In some of these examples, the native (target) motif structure is found to be absent because the nucleotides in the motif sequence either preferentially form base pairs with nucleotides in the context sequence, form a completely different structure within the motif sequence (as shown in, e.g., Fig. 2d), or simply remain unpaired (i.e., singlestranded) in a large percentage of the structures within the thermal ensemble. Such cases are typically reflected in high values of ensemble defect, which indicate large differences from the target motif structure, and in high values of Shannon entropy, which show that (parts of) the motif sequence in the thermal ensemble do not assume a single well-defined structure.

However, there are also cases in which the native motif structure is either absent or present at a low percentage, but with alternative structure(s) in the thermal ensemble very close to it. This opens up a question of the definition of the native (target) motif structure to which the structures from the thermal ensemble are compared—if the target structure is (for whatever reason) not accurate to start with, this will be reflected in the structure not being detected in the thermal ensemble. Similarly, if the structures in the ensemble fluctuate around the target structure, the results will be similar, as the structural measures take into account only the possibility of a single target structure.

As an example, we take a look at the motif SL+4, which we have defined to be 16 nt long with a native structure of ((((((….)))))) in the dot-bracket notation, that is, a 4-nt-long loop enclosed by a 6-bp-long stem (Fig. S1 and Table S1). When the motif sequence is folded on its own, ViennaRNA and oxRNA predict 𝒫= 41% and 𝒫= 76% for the target structure to occur, respectively. Similar values are for instance obtained when the motif is folded in the presence of a poly-U flanking sequence of length *L* = 30 nt (Table S2). However, if we consider in addition two alternative forms of the native motif—a shorter motif with a 5-bp-long stem and a motif with a 4-bp-long stem and an expanded, 6-nt-long loop—to be equally valid and representative of the target motif structure, we obtain values of 𝒫= 96% (ViennaRNA) and 𝒫= 93% (oxRNA) when the motif sequence is folded on its own (Table I). Similar increase is observed in the presence of a poly-U flanking sequence. These results clearly illustrate that if the target structure of a motif is identified incorrectly or if there are several similar—and equally valid—structures available to the motif, this will be reflected in the structural measures which are based on a single target structure.

**Table 1.**
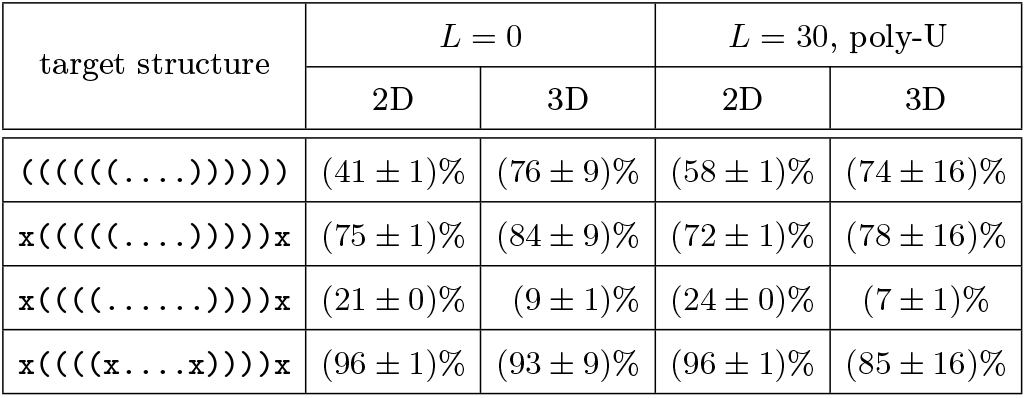
Structure probability *P* of the SL+4 motif with sequence UGACAAAUCCUUGUCA and different target motif structures. Target structures are represented in the dot-bracket notation, with x denoting “any” structure. The probabilities are listed for the motif sequence folded on its own (*L* = 0) and in the presence of a poly-U flanking sequence of length *L* = 30 nt, with predictions both at the level of secondary (2D) and tertiary (3D) structure.

### Motif sequences generally lead to lower free energies in genomic context

From the motif sequences alone (Table S1), it is difficult to predict whether or not they lead to an overall stable motif structure or to a structure which requires correct context to form. To explore if nucleotide composition of different motif sequences is responsible for the stability of their resulting structures, we have compared the free energies of secondary structure formation for the motif sequences and their (mono- and dinucleotide) shuffled versions (Fig. S4 in Supplementary Material). When motif sequences are folded on their own, most of them lead to lower free energies compared to their shuffled versions (Fig. S4a and Table S3 in Supplementary Material). Notable exceptions are motifs SL-3, SL+1, IX, and SL+5 with free energies around ∼ 0 *k*_*B*_*T* for both unshuffled and shuffled motif sequences. Despite this, we note that the motifs SL-3 and IX can be found in the structural ensemble at around ∼ 50% probability (Table S2); we have also observed that the structure of these two motifs is stabilized in the context of MS2 gRNA (Fig. S3). From the free energy perspective, adding *L* = 30 nt of genomic context on both sides of the motif preferentially stabilizes the motif sequence in comparison to its shuffled versions (Fig. S4b and Table S4 in Supplementary Material), as the ensemble free energies are, on average, at least several *k*_*B*_*T* lower for the unshuffled sequences. A notable exception is again the motif SL+1. We also observe that the motifs which we found are stable regardless of their context, such as the TR motif, do not particularly distinguish themselves from other motifs in this respect.

## DISCUSSION AND CONCLUSIONS

We have studied the structural stability of 14 hairpin motifs located in an ∼ 800-nt-long region of the RNA genome of MS2 bacteriophage. Some of these hairpin motifs have been shown to be functionally important as PS motifs, which interact with coat proteins and are thus indispensable for virus assembly [31, 32, 37]. Placing the motifs in various contexts—including their genomic sequence, poly-U sequence, and random sequences—we used different measures of structural stability on their predicted secondary and tertiary structures to determine how their structure is influenced by the flanking sequences. We found that the flanking sequence of a motif can drastically influence the structure it will assume (Fig. S3). However, the extent of this depends both on the nature of the flanking sequence as well as on the motif sequence itself. While some motifs were found to be stable no matter the context they were placed in (Fig. 4), several motifs were found to fold into the native structure only in the presence of the MS2 gRNA (Fig. 3). A motif whose structural stability stands out in particular is the TR PS, a key part of the MS2 virus assembly process [28, 30]. The structure of this motif was found to be stable even in the context of a vast majority of random sequences that were put around it, which is unlike any other motif we studied.

TR motif is also an illustrative case which shows that different structure prediction methods can lead to different results. As Fig. 5 shows, in this case, the results predicted by ViennaRNA (secondary structure) and oxRNA (tertiary structure) differ in their prediction of two base pairs. ViennaRNA-predicted secondary structure of the motif (Fig. 5a) excellently matches the native structure with a single-nucleotide bulge and a 4-nt hairpin loop. In contrast, the most common (secondary) structure of the TR motif we observe in oxRNA predictions (Fig. 5b) involves a symmetric 1-nt interior loop and a 5-nt hairpin loop. The reason for this difference is, at present unknown; however, we note that the nucleotides most important for the functionality of the TR motif—the unpaired A nucleotide in position 6 in the interior bulge and the unpaired U and A nucleotides in positions 11 and 12 in the hairpin loop [32, 78]—are present in both predictions. While the extraction of secondary structure from tertiary structure can be an imprecise way of structure comparison, inspection of per-nucleotide energies of hydrogen bonding for the two tertiary structures indicate that this is not the case for the prediction of the TR motif structure (panels (d) and (e) of Fig. 5). In general, validation of predicted structures of both TR and other motifs would require experimental input using, for instance, probing experiments, which can also be applied to structural ensembles [79].

**Figure 5.**
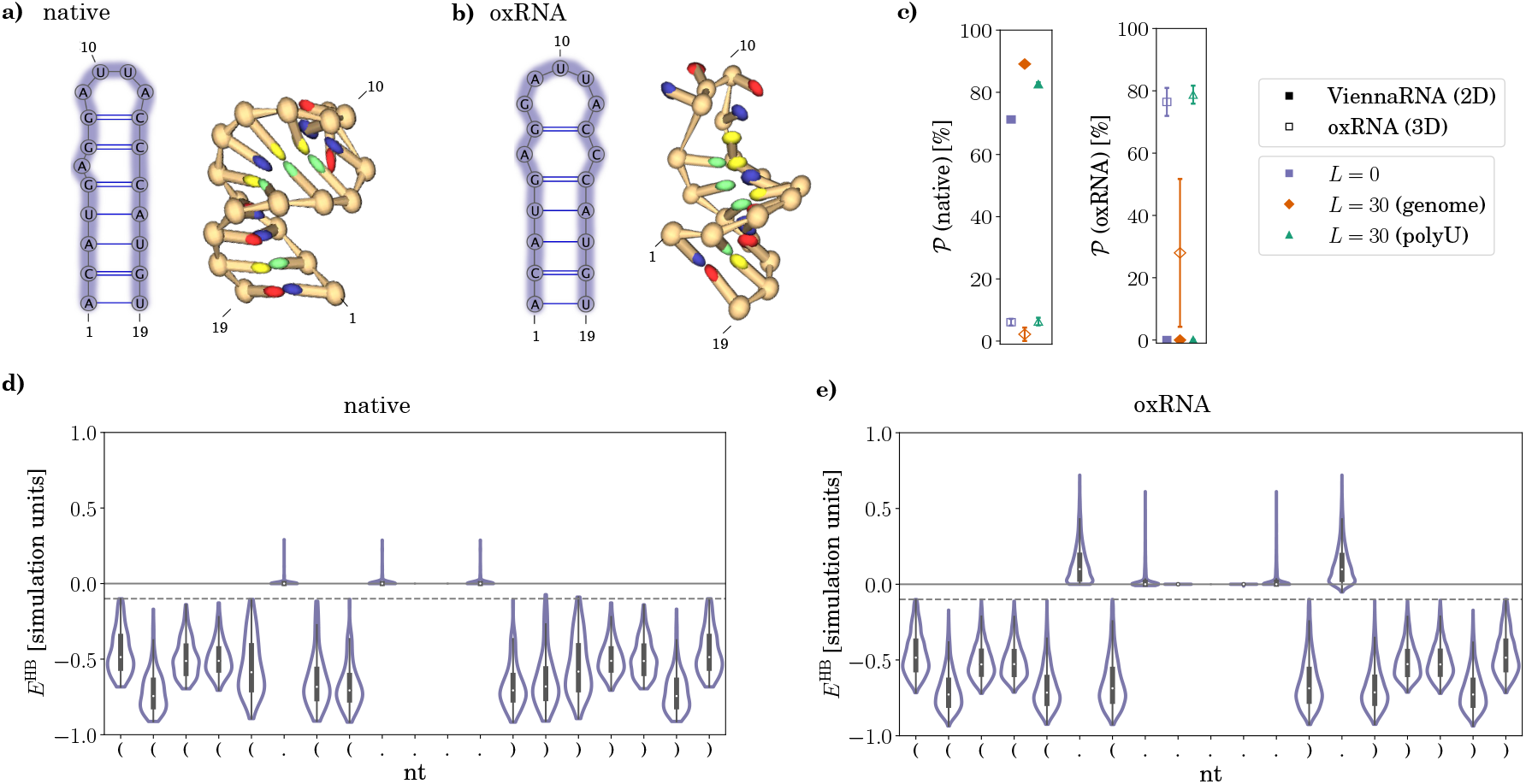
Predicted structures of the TR motif. **(a)** Secondary and tertiary structure of the native TR motif, which is also the most probable structure predicted by ViennaRNA in different contexts. **(b)** Secondary and tertiary structure of the most common TR motif structure as predicted by oxRNA, occurring with a high probability in most contexts. **(c)** Structure probability of the ViennaRNA(panel (a)) and oxRNA-predicted (panel (b)) structures of the TR motif in different contexts. **(d)**–**(e)** Violin plots of per-nucleotide hydrogen bonding energies for the motifs in panels (a) and (b), respectively, obtained from a single oxRNA Lrun. For each nucleotide, the sum of its hydrogen bonding energies with all other nucleotides in the motif is shown, i.e., 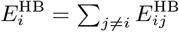. Dashed lines show the cut-off for a base pair to be considered formed (i.e., 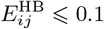 in simulation units).

There are other aspects of RNA folding that can influence the study of their structural stability. For instance, it is likely that the structure of some PS motifs is stabilized by their interaction with the MS2 coat proteins [32]. Since our study focused solely on the changes in the motif structure in the presence of flanking RNA sequences, any conformational changes or stabilizing effects incurred by interactions with the environment have been neglected. Similarly, cotranscriptional folding is another factor whose importance for RNA folding is becoming increasingly more recognised [80]. Given the fairly short nature of the RNA sequences that we have examined we believe that cotranscriptional folding does not play a significant role; this is further supported by the experimental determinations of RNA structure of the MS2 genome in the region that we studied and on which the native motif structures we use are based. Lastly, in light of the relatively high density of hairpin motifs in the studied region of MS2 gRNA (the 14 motifs represent approximately one third of the region), it is also likely that the structure formation of *neighbouring* motifs is not independent, and that tertiary interactions between different motifs can stabilize their individual structure. This represents an interesting direction for future studies, which will have to take into account tertiary interactions and three-dimensional structure formation of a fairly large region.

We have also observed that the reliability of the different structural measures we have employed depends on how well the target structure of the motif is known. We have identified cases where a slight change in the target motif structure (for instance, shortening of the stem or expansion of the hairpin loop) led to a significant increase in the presence of the correct motif structure in different context (Table I). The choice of a single target structure can thus influence the results obtained through different structural measures if the motif structure is not precisely known or if the motif assumes several closely-related structures in the thermal ensemble. However, our approach can be easily extended to such cases by implementing inexact structure matching in the structural measures, allowing for select nucleotides to participate in different structures without incurring a penalty.

Our study provides new insights into the observations that context—flanking sequence up- and down-stream of an RNA sequence—can influence the formation of RNA structure, showing that the structure of an RNA motif can be very sensitive to its context. The structural stability of a motif can, in turn, provide insight into its function—the TR motif, the highest affinity PS in the MS2 gRNA, is also by far the most structurally stable in different contexts. This is particularly relevant for +ssRNA bacteriophages, which are incredibly diverse [51] and exhibit differences in distribution of PSs [52] and their assembly processes [53] despite being closely related. In light of the numerous functional RNA structures that occur in the midst of larger RNA molecules, the importance of our findings reaches beyond the specific hairpin motifs in the MS2 gRNA and should be of interest also in studies of RBP-binding motifs, mitochondrial tRNA structures, and bioengineered RNAs utilizing known RNA motifs, including PSs found in +ssRNA viruses.

## ACKNOWLEDGMENTS

We acknowledge support by Slovenian Research Agency (ARRS) under contract no. P1-0055.

## AUTHOR CONTRIBUTIONS

A.B. designed the research. V.B. performed the research. V.B. and A.B. analysed the results. A.B. wrote the manuscript and V.B. and A.B. revised the manuscript.

## DATA AVAILABILITY

The data that support the findings of this study are available in OSF with the identifier [DOI 10.17605/OSF.IO/37H8Z].

## References

[1] L. Statello, C.-J. Guo, L.-L. Chen, and M. Huarte, Nat. Rev. Mol. Cell Biol. 22, 96 (2021).

[2] F. Michelini, A. P. Jalihal, S. Francia, C. Meers, Z. T. Neeb, F. Rossiello, U. Gioia, J. Aguado, C. Jones-Weinert, B. Luke, et al., Chem. Rev. 118, 4365 (2018).

[3] E. C. Holmes, The evolution and emergence of RNA viruses (Oxford University Press, 2009).

[4] X.-W. Wang, C.-X. Liu, L.-L. Chen, and Q. C. Zhang, Nat. Chem. Biol. 17, 755 (2021).

[5] J. Gorodkin and W. L. Ruzzo, RNA sequence, structure, and function: Computational and bioinformatic methods (Springer, 2014).

[6] S. A. Mortimer, M. A. Kidwell, and J. A. Doudna, Nat. Rev. Genet. 15, 469 (2014).

[7] P. Brion and E. Westhof, Annu. Rev. Biophys. Biomol. Struct. 26, 113 (1997).

[8] L. R. Ganser, M. L. Kelly, D. Herschlag, and H. M. Al-Hashimi, Nat. Rev. Mol. Cell Biol. 20, 474 (2019).

[9] A. M. Mustoe, C. L. Brooks, and H. M. Al-Hashimi, Annu. Rev. Biochem. 83, 441 (2014).

[10] A. Schneemann, Annu. Rev. Microbiol. 60, 51 (2006).

[11] N. Patel, E. C. Dykeman, R. H. Coutts, G. P. Lomonossoff, D. J. Rowlands, S. E. Phillips, N. Ranson, R. Twarock, R. Tuma, and P. G. Stockley, Proc. Natl. Acad. Sci. USA 112, 2227 (2015).

[12] P. Simmonds, A. Tuplin, and D. J. Evans, RNA 10, 1337 (2004).

[13] A. M. Yoffe, P. Prinsen, A. Gopal, C. M. Knobler, W. M. Gelbart, and A. Ben-Shaul, Proc. Natl. Acad. Sci. USA 105, 16153 (2008).

[14] L. Tubiana, A. Božič, C. Micheletti, and R. Podgornik, Biophys. J. 108, 194 (2015).

[15] R. Twarock, G. J. Towers, and P. G. Stockley, Trends Microbiol. 32, 17 (2024).

[16] N. B. Leontis and E. Westhof, Curr. Op. Struct. Biol. 13, 300 (2003).

[17] N. B. Leontis, A. Lescoute, and E. Westhof, Curr. Op. Struct. Biol. 16, 279 (2006).

[18] P. Svoboda and A. D. Cara, Cell. Mol. Life Sci. 63, 901 (2006).

[19] L. Ye, U. B. Ambi, M. Olguin-Nava, A.-S. Gribling-Burrer, S. Ahmad, P. Bohn, M. M. Weber, and R. P. Smyth, Viruses 13, 1788 (2021).

[20] M. Comas-Garcia, Viruses 11, 253 (2019).

[21] S. Tetter, N. Terasaka, A. Steinauer, R. J. Bingham, S. Clark, A. J. Scott, N. Patel, M. Leibundgut, E. Wroblewski, N. Ban, et al., Science 372, 1220 (2021).

[22] K. V. Gorzelnik, Z. Cui, C. A. Reed, J. Jakana, R. Young, and J. Zhang, Proc. Natl. Acad. Sci. USA 113, 11519 (2016).

[23] N. Patel, E. Wroblewski, G. Leonov, S. E. Phillips, R. Tuma, R. Twarock, and P. G. Stockley, Proc. Natl. Acad. Sci. USA 114, 12255 (2017).

[24] K. E. Watters, K. Choudhary, S. Aviran, J. B. Lucks, K. L. Perry, and J. R. Thompson, Nucleic Acids Res. 46, 2573 (2018).

[25] R. F. Garmann, A. M. Goldfain, C. R. Tanimoto, C. E. Beren, F. F. Vasquez, D. A. Villarreal, C. M. Knobler, W. M. Gelbart, and V. N. Manoharan, Proc. Natl. Acad. Sci. USA 119, e2206292119 (2022).

[26] L. A. Williams, A. Neophytou, R. F. Garmann, D. Chakrabarti, and V. N. Manoharan, Nanoscale 16, 3121 (2024).

[27] P. G. Stockley, R. Twarock, S. E. Bakker, A. M. Barker, A. Borodavka, E. Dykeman, R. J. Ford, A. R. Pearson, S. E. Phillips, N. A. Ranson, et al., J. Biol. Phys. 39, 277 (2013).

[28] R. Twarock, R. J. Bingham, E. C. Dykeman, and P. G. Stockley, Curr. Op. Virol. 31, 74 (2018).

[29] K. Toropova, G. Basnak, R. Twarock, P. G. Stockley, and N. A. Ranson, J. Mol. Biol. 375, 824 (2008).

[30] E. C. Dykeman, P. G. Stockley, and R. Twarock, J. Mol. Biol. 425, 3235 (2013).

[31] Ó. Rolfsson, S. Middleton, I. W. Manfield, S. J. White, B. Fan, R. Vaughan, N. A. Ranson, E. Dykeman, R. Twarock, J. Ford, et al., J. Mol. Biol. 428, 431 (2016).

[32] X. Dai, Z. Li, M. Lai, S. Shu, Y. Du, Z. H. Zhou, and R. Sun, Nature 541, 112 (2017).

[33] R. I. Koning, J. Gomez-Blanco, I. Akopjana, J. Vargas, A. Kazaks, K. Tars, J. M. Carazo, and A. J. Koster, Nat. Commun. 7, 12524 (2016).

[34] P. Pumpens, Single-stranded RNA phages: From molecular biology to nanotechnology (CRC Press, 2020).

[35] V. S. Farafonov, M. Stich, and D. A. Nerukh, J. Chem. Theory Comput. 19, 7924 (2023).

[36] J. Rudnick and R. Bruinsma, Phys. Rev. Lett. 94, 038101 (2005).

[37] R. Twarock, G. Leonov, and P. G. Stockley, Nat. Commun. 9, 2021 (2018).

[38] J. D. Farrell, J. Dobnikar, and R. Podgornik, Phys. Rev. Res. 5, L012040 (2023).

[39] U. Ohler, S. Yekta, L. P. Lim, D. P. Bartel, and C. B. Burge, RNA 10, 1309 (2004).

[40] Y. Zeng and B. R. Cullen, J. Biol. Chem. 280, 27595 (2005).

[41] J. M. Taliaferro, N. J. Lambert, P. H. Sudmant, D. Dominguez, J. J. Merkin, M. S. Alexis, C. A. Bazile, and C. B. Burge, Mol. Cell 64, 294 (2016).

[42] D. Dominguez, P. Freese, M. S. Alexis, A. Su, M. Hochman, T. Palden, C. Bazile, N. J. Lambert, E. L. Van Nostrand, G. A. Pratt, et al., Mol. Cell 70, 854 (2018).

[43] T. C. Lan, M. F. Allan, L. E. Malsick, J. Z. Woo, C. Zhu, F. Zhang, S. Khandwala, S. S. Nyeo, Y. Sun, J. U. Guo, et al., Nat. Commun. 13, 1128 (2022).

[44] Y. S. Jeong, J. F. Repass, Y.-N. Kim, S.-M. Hwang, and S. Makino, Virology 217, 311 (1996).

[45] Y. Wang, Z. Wang, T. Liu, S. Gong, and W. Zhang, RNA 24, 1229 (2018).

[46] X. Sun and A. E. Simon, J. Virol. 77, 7880 (2003).

[47] S. Busan and K. M. Weeks, Biochemistry 52, 8219 (2013).

[48] E. A. Jolley, K. M. Bormes, and P. C. Bevilacqua, J. Mol. Biol. 434, 167786 (2022).

[49] S. Cardinale and A. P. Arkin, Biotech. J. 7, 856 (2012).

[50] C. Ling, P. Hung, and L. Overby, Virology 40, 920 (1970).

[51] K. Tars, ssRNA phages: Life cycle, structure and applications, in Biocommunication of Phages, edited by G. Witzany (Springer International Publishing, Cham, 2020) pp. 261–292.

[52] J. Thongchol, Z. Lill, Z. Hoover, and J. Zhang, Viruses 15, 1985 (2023).

[53] S. Hamilton, T. Modi, P. Šulc, and B. Ozkan, bioRxiv, 2023 (2023).

[54] P. Radecki, R. Uppuluri, K. Deshpande, and S. Aviran, RNA Biol. 18, 521 (2021).

[55] D. Ojala, J. Montoya, and G. Attardi, Nature 290, 470 (1981).

[56] T. Suzuki, A. Nagao, and T. Suzuki, Annu. Rev. Genet. 45, 299 (2011).

[57] L. M. Wittenhagen and S. O. Kelley, Trends Biochem. Sci. 28, 605 (2003).

[58] R. Kent, S. Halliwell, K. Young, N. Swainston, and N. Dixon, ACS Synth. Biol. 7, 1660 (2018).

[59] N. Katz, R. Cohen, O. Solomon, B. Kaufmann, O. Atar, Z. Yakhini, S. Goldberg, and R. Amit, ACS Synth. Biol. 7, 2765 (2018).

[60] N. Katz, E. Tripto, N. Granik, S. Goldberg, O. Atar, Z. Yakhini, Y. Orenstein, and R. Amit, Nat. Commun. 12, 1576 (2021).

[61] N. Granik, N. Katz, O. Willinger, S. Goldberg, and R. Amit, Nat. Commun. 13, 6811 (2022).

[62] R. Lorenz, S. H. Bernhart, C. Höner zu Siederdissen, H. Tafer, C. Flamm, P. F. Stadler, and I. L. Hofacker, Algorithms Mol. Biol. 6, 1 (2011).

[63] T. Schlick and A. M. Pyle, Biophys. J. 113, 225 (2017).

[64] C. T. Woods, L. Lackey, B. Williams, N. V. Dokholyan, D. Gotz, and A. Laederach, Biophys. J. 113, 290 (2017).

[65] P. Šulc, F. Romano, T. E. Ouldridge, L. Rovigatti, J. P. K. Doye, and A. A. Louis, J. Chem. Phys. 137, 135101 (2012).

[66] P. Šulc, F. Romano, T. E. Ouldridge, J. P. K. Doye, and A. A. Louis, J. Chem. Phys. 140, 235102 (2014).

[67] M. L. Sample, M. Matthies, and P. Šulc, in 2023 Winter Simulation Conference (WSC) (IEEE, 2023) pp. 91–105.

[68] T. E. Ouldridge, A. A. Louis, and J. P. Doye, J. Chem. Phys. 134 (2011).

[69] J. S. Martin, Entropy 16, 1331 (2014).

[70] M. Ward, E. Courtney, and E. Rivas, Nucleic Acids Res. 51, e40 (2023).

[71] R. M. Dirks, M. Lin, E. Winfree, and N. A. Pierce, Nucleic Acids Res. 32, 1392 (2004).

[72] N. A. Siegfried, S. Busan, G. M. Rice, J. A. Nelson, and K. M. Weeks, Nat. Methods 11, 959 (2014).

[73] I. Manfredonia, C. Nithin, A. Ponce-Salvatierra, P. Ghosh, T. K. Wirecki, T. Marinus, N. S. Ogando, E. J. Snijder, M. J. van Hemert, J. M. Bujnicki, et al., Nucleic Acids Res. 48, 12436 (2020).

[74] N. C. Huston, H. Wan, M. S. Strine, R. d. C. A. Tavares, C. B. Wilen, and A. M. Pyle, Mol. Cell 81, 584 (2021).

[75] F. Pedregosa, G. Varoquaux, A. Gramfort, V. Michel, B. Thirion, O. Grisel, M. Blondel, P. Prettenhofer, R. Weiss, V. Dubourg, J. Vanderplas, A. Passos, D. Cournapeau, M. Brucher, M. Perrot, and E. Duchesnay, J. Mach. Learn. Res. 12, 2825 (2011).

[76] G. J. McLachlan and S. Rathnayake, WIREs Data Min. Knowl. Discov. 4, 341 (2014).

[77] M. Jiang, J. Anderson, J. Gillespie, and M. Mayne, BMC Bioinf. 9, 1 (2008).

[78] K. Valegård, J. B. Murray, N. J. Stonehouse, S. van den Worm, P. G. Stockley, and L. Liljas, J. Mol. Biol. 270, 724 (1997).

[79] R. C. Spitale and D. Incarnato, Nat. Rev. Genet. 24, 178 (2023).

[80] D. Z. Bushhouse, E. K. Choi, L. M. Hertz, and J. B. Lucks, J. Mol. Biol. 434, 167665 (2022).

