## Supplementary Material for "Context-dependent structure formation of hairpin motifs in bacteriophage MS2 genomic RNA"

Veronika Bukina
*Department of Theoretical Physics, Jožef Stefan Institute, Ljubljana, Slovenia and*
*Department of Physics, Faculty of Mathematics and Physics, University of Ljubljana, Ljubljana, Slovenia*

Anže Božič*
*Department of Theoretical Physics, Jožef Stefan Institute, Ljubljana, Slovenia*

*

#### I. HAIRPIN MOTIFS IN THE MS2 GENOME

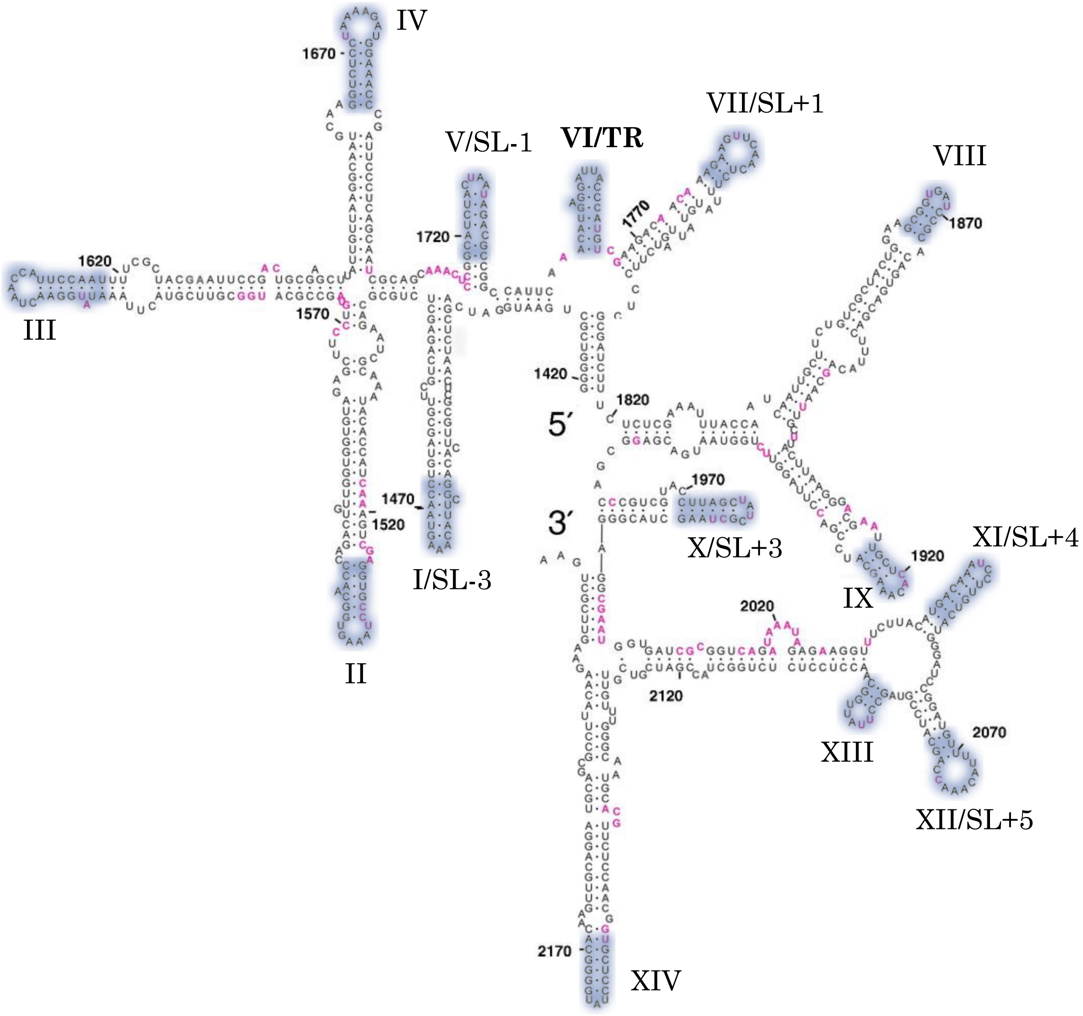


Figure S1. Secondary structure of the [1419–2211] nt region of the MS2 genome, which includes 14 hairpin motifs analyzed in our study. Motifs are highlighted in violet colour, their labels following the notation introduced by Rolfsson *et al.* [1]. Image modified from Ref. [1] under the Creative Commons CC-BY license.

| motif[a] | sequence–structure | position[b] | length |
|---|---|---|---|
| I/SL-3 | GGCUUACAAAGUAACC<br>((.((((...)))))) | 1457 | 16 nt |
| II | GGUGCCUAAAGUGGCAACC<br>((((((......))).))) | 1526 | 19 nt |
| III | AUAUGGAACUAACCAUUCCAAU<br>((.(((((.......))))))) | 1597 | 22 nt |
| IV | GGUCUCCUAAAAGAUGGAAACC<br>(((.(((........))).))) | 1665 | 22 nt |
| V/SL-1 | GCAUCUACUAAUAGACGC<br>((.((((....)))).)) | 1717 | 18 nt |
| VI/TR | ACAUGAGGAUUACCCAUGU<br>(((((.((....))))))) | 1746 | 19 nt |
| VII/SL+1 | AGAAGUUCAACUCU<br>(((........))) | 1777 | 14 nt |
| VIII | GCGGUGAUCCGC<br>((((....)))) | 1861 | 12 nt |
| IX | GCUCACAAAGC<br>(((.....))) | 1916 | 11 nt |
| X/SL+3 | CUUAGCUAUCGCUAAG<br>((((((....)))))) | 1970 | 16 nt |
| XI/SL+4 | UGACAAAUCCUUGUCA<br>((((((....)))))) | 2039 | 16 nt |
| XII/SL+5 | GUUUUACAAACCAGC<br>(((.........))) | 2067 | 15 nt |
| XIII | GCCUUAUUGGC<br>(((.....))) | 2089 | 11 nt |
| XIV | UGCUCCUAUGGGGCA<br>((((((...)))))) | 2156 | 15 nt |

[a] Motif name (Rolfsson *et al.* [1]).
[b] Starting position of the motif in the MS2 genome (with genome start at position 1).

Table S1. Sequences and structures of the 14 MS2 hairpin motifs analysed in this work. The table also lists the length of the motif and its starting position within the MS2 genome.

#### II. GAUSSIAN MIXTURE MODELS

As described in the Methods in the main text, adding 500 different random flanking sequences to a motif sequence results in a diverse ensemble of motif secondary structures. In order to represent the overall structural properties of a motif in the presence of random flanking sequences in the simplest fashion, we apply a Gaussian mixture model [2] to the distributions of the structural measures in order to separate, at least approximately, the contributions of different populations of motif structures. Typically, mixtures of two or three components suffice to describe the ensemble, as determined by the minimum of the BIC curve [3]. In order to keep the comparison simple, we limit ourselves to at most three components.

Figure S2 illustrates this approach on the example of the motif SL-3. Each of the panels (b)–(d) shows a histogram of the native motif structure probability $\mathcal{P}$ for 500 random flanking sequences of different lenghts $L$. The panels also show the two- and three-component Gaussian mixture distributions together with their corresponding averages. Each of the components is also assigned a weight depending on the proportion of random sequences that fall under it.

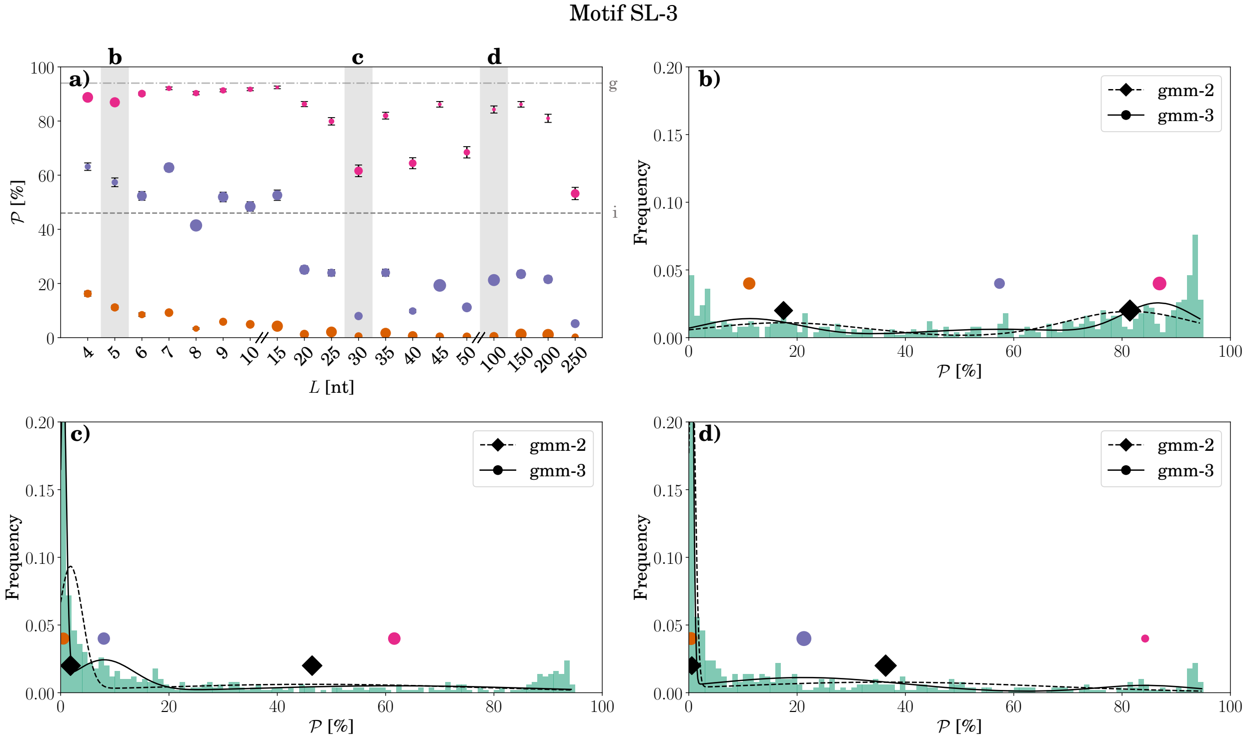


Figure S2. Gaussian mixture model shown on the example of structure probability $\mathcal{P}(\mathcal{M})$ of the motif SL-3 (Table S1). **(a)** Structure probability as a function of the flanking sequence length $L$ in presence of the MS2 genome. The distribution for the $\mathcal{P}(\mathcal{M})$ at each value of $L$ consists of 500 random sequences. To this distribution, we apply a Gaussian mixture model with either 2 or 3 components and select the one with the lowest BIC value. The size of each point corresponds to the weight of the mixture component, and its colour denotes its order in the mixture (first, second, or third, where applicable). Error bars correspond to the standard deviation of the distribution. Dashed and dot-dashed lines correspond to the values obtained for an isolated motif sequence (i) or motif sequence in the presence of the MS2 genome (g), respectively. **(b)**–**(d)** Histograms of the distributions of $\mathcal{P}(\mathcal{M})$ at three different values of flanking sequence length $L = 5$, 30, and 100 (also marked in panel (a)). For each of the three distributions, the Gaussian mixture density with 2 or 3 components (lines) and their corresponding means (symbols) are shown. The size of each symbol again corresponds to the weight of the mixture component.

### III. STRUCTURAL MEASURES OF THE MS2 HAIRPIN MOTIFS

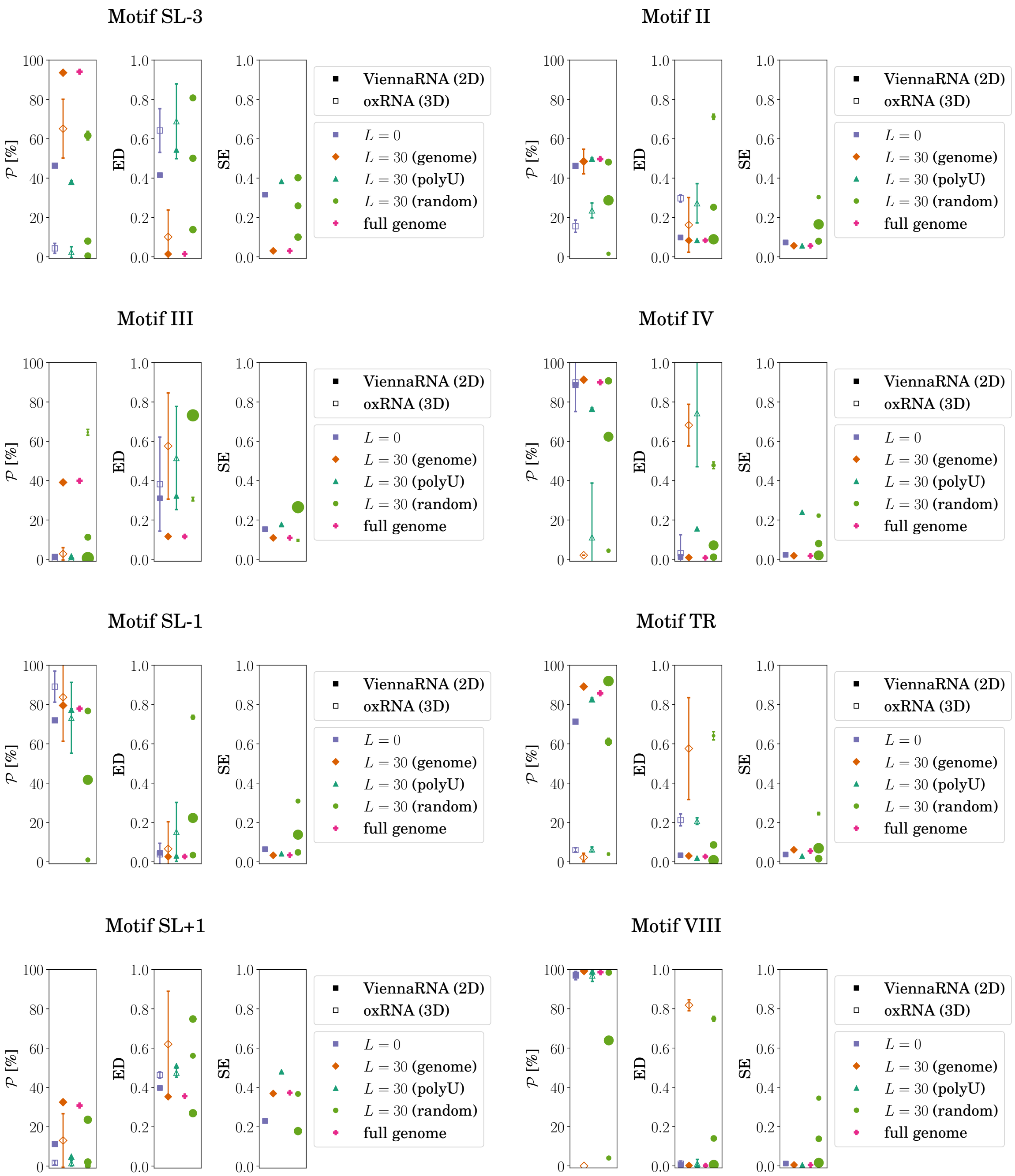


Figure S3. (Caption on next page.)

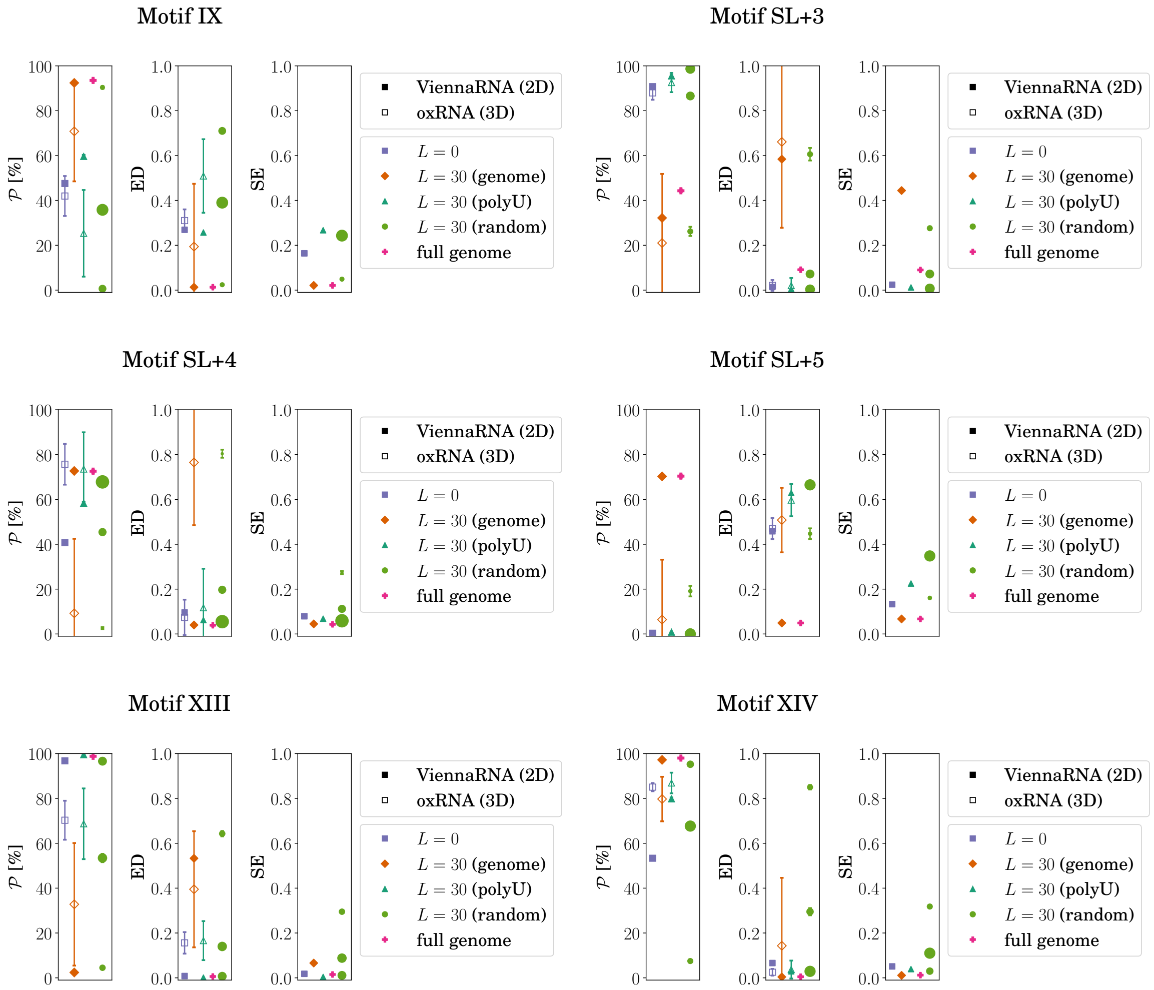


Figure S3. *(cont'd)* Structure probability, ensemble defect, and Shannon entropy of all 14 hairpin motif structures as determined by ViennaRNA (full symbols) and oxRNA (empty symbols). The values are shown for the motif in absence of any flanking sequences ($L = 0$) as well as in the presence of different flanking sequences—genomic, poly-U, and random—of length $L = 30$ nt. For the predicted secondary structure, the values are also shown for the motif in the context of the full gRNA. Size of the symbols for random flanking sequences corresponds to the weight of the corresponding component of the Gaussian mixture model (see Methods and Figure S2).

| motif | $L = 0$ | | $L = 30$ | | | | | | | full genome |
|---|---|---|---|---|---|---|---|---|---|---|
| | | | genome | | poly-U | | random | | | |
| | 2D | 3D | 2D | 3D | 2D | 3D | 2D | | | 2D |
| I/SL-3 | $0.46 \pm 0.01$ | $0.04 \pm 0.03$ | $0.94 \pm 0.0$ | $0.65 \pm 0.15$ | $0.38 \pm 0.01$ | $0.02 \pm 0.03$ | $0.62 \pm 0.02$ (34%) | $0.08 \pm 0.01$ (34%) | $0.01 \pm 0.0$ (32%) | $0.94 \pm 0.0$ |
| II | $0.46 \pm 0.01$ | $0.15 \pm 0.03$ | $0.49 \pm 0.01$ | $0.49 \pm 0.06$ | $0.50 \pm 0.01$ | $0.24 \pm 0.04$ | $0.48 \pm 0.01$ (32%) | $0.29 \pm 0.01$ (49%) | $0.02 \pm 0.0$ (19%) | $0.50 \pm 0.01$ |
| III | $0.01 \pm 0.0$ | 0.0 | $0.40 \pm 0.01$ | $0.03 \pm 0.03$ | $0.02 \pm 0.0$ | $0.01 \pm 0.0$ | $0.65 \pm 0.02$ (9%) | $0.01 \pm 0.12$ (32%) | $0.01 \pm 0.0$ (59%) | $0.40 \pm 0.01$ |
| IV | $0.89 \pm 0.01$ | $0.90 \pm 0.15$ | $0.91 \pm 0.01$ | $0.02 \pm 0.0$ | $0.76 \pm 0.01$ | $0.11 \pm 0.28$ | $0.91 \pm 0.01$ (34%) | $0.62 \pm 0.02$ (47%) | $0.04 \pm 0.01$ (19%) | $0.90 \pm 0.01$ |
| V/SL-1 | $0.72 \pm 0.01$ | $0.89 \pm 0.08$ | $0.80 \pm 0.01$ | $0.84 \pm 0.23$ | $0.77 \pm 0.01$ | $0.73 \pm 0.18$ | $0.77 \pm 0.01$ (30%) | $0.42 \pm 0.02$ (48%) | $0.01 \pm 0.0$ (22%) | $0.78 \pm 0.01$ |
| VI/TR | $0.71 \pm 0.01$ | $0.06 \pm 0.01$ | $0.89 \pm 0.01$ | $0.02 \pm 0.02$ | $0.83 \pm 0.01$ | $0.06 \pm 0.01$ | $0.92 \pm 0.01$ (50%) | $0.61 \pm 0.02$ (35%) | $0.04 \pm 0.01$ (15%) | $0.86 \pm 0.01$ |
| VII/SL+1 | $0.11 \pm 0.0$ | $0.02 \pm 0.01$ | $0.31 \pm 0.01$ | $0.13 \pm 0.14$ | $0.05 \pm 0.0$ | $0.01 \pm 0.01$ | $0.24 \pm 0.01$ (39%) | $0.02 \pm 0.0$ (35%) | 0.0 (26%) | $0.31 \pm 0.01$ |
| VIII | $0.97 \pm 0.0$ | $0.97 \pm 0.02$ | $0.99 \pm 0.0$ | 0.0 | $0.99 \pm 0.0$ | $0.97 \pm 0.03$ | $0.98 \pm 0.0$ (30%) | $0.649 \pm 0.02$ (47%) | $0.04 \pm 0.0$ (23%) | $0.99 \pm 0.0$ |
| IX | $0.48 \pm 0.01$ | $0.42 \pm 0.09$ | $0.92 \pm 0.01$ | $0.71 \pm 0.22$ | $0.60 \pm 0.01$ | $0.25 \pm 0.19$ | $0.90 \pm 0.01$ (19%) | $0.36 \pm 0.02$ (50%) | $0.01 \pm 0.0$ (32%) | $0.94 \pm 0.0$ |
| X/SL+3 | $0.91 \pm 0.01$ | $0.88 \pm 0.03$ | $0.32 \pm 0.01$ | $0.21 \pm 0.31$ | $0.96 \pm 0.0$ | $0.93 \pm 0.04$ | $0.99 \pm 0.0$ (41%) | $0.87 \pm 0.01$ (36%) | $0.26 \pm 0.02$ (23%) | $0.44 \pm 0.01$ |
| XI/SL+4 | $0.41 \pm 0.01$ | $0.76 \pm 0.09$ | $0.73 \pm 0.01$ | $0.09 \pm 0.33$ | $0.58 \pm 0.01$ | $0.74 \pm 0.16$ | $0.68 \pm 0.01$ (57%) | $0.45 \pm 0.01$ (33%) | $0.03 \pm 0.0$ (10%) | $0.73 \pm 0.01$ |
| XII/SL+5 | 0.0 | 0.0 | $0.70 \pm 0.01$ | $0.06 \pm 0.27$ | $0.01 \pm 0.0$ | 0.0 | $0.19 \pm 0.02$ (16%) | $0.01 \pm 0.0$ (38%) | 0.0 (47%) | $0.70 \pm 0.01$ |
| XIII | $0.97 \pm 0.0$ | $0.70 \pm 0.09$ | $0.02 \pm 0.0$ | $0.33 \pm 0.27$ | $0.99 \pm 0.0$ | $0.69 \pm 0.16$ | $0.97 \pm 0.0$ (36%) | $0.53 \pm 0.02$ (39%) | $0.05 \pm 0.0$ (25%) | $0.99 \pm 0.0$ |
| XIV | $0.53 \pm 0.01$ | $0.85 \pm 0.02$ | $0.97 \pm 0.0$ | $0.80 \pm 0.10$ | $0.80 \pm 0.01$ | $0.87 \pm 0.05$ | $0.95 \pm 0.0$ (30%) | $0.68 \pm 0.01$ (47%) | $0.08 \pm 0.01$ (24%) | $0.98 \pm 0.0$ |

Table S2. Structure probability $\mathcal{P}$ of the 14 hairpin motifs in different contexts: when the motif sequence if folded alone ($L = 0$), in the presence of the full genome, or in the presence of different flanking sequences of length $L = 30$ nt (either taken from the MS2 gRNA, consisting solely of U bases, or generated randomly). For the random flanking sequences, averages and errorbars of two or three components of a Gaussian mixture model are given, together with the weight of each component in parentheses (cf. also Fig. S2). The structures were predicted either at the level of secondary structure (2D) using ViennaRNA or at the level of tertiary structure (3D) using oxRNA, from which the secondary structure was then extracted (see also Methods in the main text). Statistical error for the 2D case is estimated according to multinomial distribution due to finite number of structures. The total error for $\mathcal{P}$ for the case of random flanking sequences additionally includes model error due to the limited number of simulations ($N = 500$). The error for the 3D case is the corrected sample standard deviation over 20 independent simulation runs.

#### IV. FOLDING ENERGIES OF MOTIF STRUCTURAL ENSEMBLES

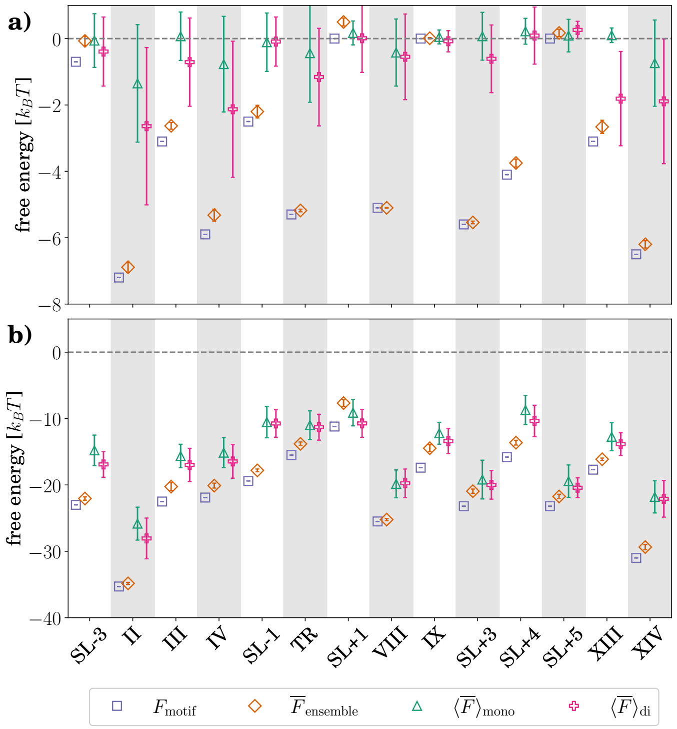


Figure S4. Minimum free energy of the motif sequence $F_{\mathrm{motif}}$ and the average folding free energies of 3000 secondary structures sampled from the thermal ensemble of the motif sequence $\overline{F}_{\mathrm{ensemble}}$ and from the thermal ensembles of both monon- and dinucleotide shuffled motif sequences ($\overline{F}_{\mathrm{mono}}$ and $\overline{F}_{\mathrm{di}}$, respectively). We produce 100 unique shuffles for each case where possible; due to the short motif sequence length, the number of dinucleotide shuffles is lower for motifs IX, XIII, and SL+5 (18, 12, and 84 dinucleotide shuffles, respectively); averaging over the shuffled sequences is denoted by $\langle\cdot\rangle$. Folding free energies are shown for **(a)** motif sequences and their shuffled versions only and **(b)** motif sequences and their shuffled versions in the presence of $L = 30$ nt of MS2 gRNA context; cf. also Tables S3 and S4.

| motif | $F_{\text{motif}}$ $[k_BT]$ | $\overline{F}_{\text{ensemble}}$ $[k_BT]$ | $\langle\overline{F}\rangle_{\text{mono}}$ $[k_BT]$ | $\langle\overline{F}\rangle_{\text{di}}$ $[k_BT]$ |
|---|---|---|---|---|
| I/SL-3 | $-0.7$ | $-0.07 \pm 0.14$ | $-0.06 \pm 0.81$ | $-0.39 \pm 1.04$ |
| II | $-7.2$ | $-6.89 \pm 0.15$ | $-1.35 \pm 1.77$ | $-2.64 \pm 2.37$ |
| III | $-3.1$ | $-2.63 \pm 0.10$ | $0.07 \pm 0.73$ | $-0.71 \pm 1.33$ |
| IV | $-5.9$ | $-5.32 \pm 0.18$ | $-0.77 \pm 1.44$ | $-2.13 \pm 2.05$ |
| V/SL-1 | $-2.5$ | $-2.20 \pm 0.19$ | $-0.11 \pm 0.88$ | $-0.09 \pm 0.74$ |
| VI/TR | $-5.3$ | $-5.18 \pm 0.04$ | $-0.44 \pm 1.48$ | $-1.16 \pm 1.47$ |
| VII/SL+1 | $0.0$ | $0.50 \pm 0.12$ | $0.17 \pm 0.36$ | $0.01 \pm 1.03$ |
| VIII | $-5.1$ | $-5.10 \pm 0.01$ | $-0.42 \pm 1.01$ | $-0.55 \pm 1.29$ |
| IX | $0.0$ | $0.01 \pm 0.02$ | $0.05 \pm 0.21$ | $-0.08 \pm 0.32$ |
| X/SL+3 | $-5.6$ | $-5.54 \pm 0.04$ | $0.07 \pm 0.72$ | $-0.61 \pm 1.02$ |
| XI/SL+4 | $-4.1$ | $-3.75 \pm 0.12$ | $0.22 \pm 0.39$ | $0.09 \pm 0.86$ |
| XII/SL+5 | $0.0$ | $0.17 \pm 0.11$ | $0.09 \pm 0.49$ | $0.26 \pm 0.26$ |
| XIII | $-3.1$ | $-2.66 \pm 0.20$ | $0.10 \pm 0.22$ | $-1.81 \pm 1.42$ |
| XIV | $-6.5$ | $-6.20 \pm 0.11$ | $-0.74 \pm 1.30$ | $-1.89 \pm 1.88$ |

Table S3. Folding free energy of the 14 MS2 isolated hairpin motifs. Listed are the minimum free energy of the motif sequence $F_{\text{motif}}$, the average free energy of the structural ensemble obtained for the motif sequence $\overline{F}_{\text{ensemble}}$, and the average free energies of the structural ensemble of the mono- and dinucleotide shuffled motif sequence ($\langle\overline{F}\rangle_{\text{mono}}$ and $\langle\overline{F}\rangle_{\text{di}}$, respectively). We produce 100 unique shuffles for each case where possible; due to the short motif sequence length, the number of dinucleotide shuffles is lower for motifs IX, XIII, and SL+5 (18, 12, and 84 dinucleotide shuffles, respectively).

| motif | $F_{\text{motif}}$ $[k_BT]$ | $\overline{F}_{\text{ensemble}}$ $[k_BT]$ | $\langle\overline{F}\rangle_{\text{mono}}$ $[k_BT]$ | $\langle\overline{F}\rangle_{\text{di}}$ $[k_BT]$ |
|---|---|---|---|---|
| I/SL-3 | $-23.0$ | $-22.05 \pm 0.25$ | $-14.76 \pm 2.30$ | $-16.91 \pm 1.93$ |
| II | $-35.3$ | $-34.83 \pm 0.15$ | $-25.83 \pm 2.48$ | $-28.06 \pm 3.07$ |
| III | $-22.5$ | $-20.25 \pm 0.59$ | $-15.64 \pm 1.77$ | $-16.98 \pm 2.50$ |
| IV | $-21.9$ | $-20.10 \pm 0.37$ | $-15.14 \pm 2.25$ | $-16.46 \pm 2.52$ |
| V/SL-1 | $-19.4$ | $-17.81 \pm 0.21$ | $-10.53 \pm 2.37$ | $-10.74 \pm 2.07$ |
| VI/TR | $-15.5$ | $-13.81 \pm 0.28$ | $-10.99 \pm 2.15$ | $-11.30 \pm 1.95$ |
| VII/SL+1 | $-11.2$ | $-7.67 \pm 0.51$ | $-9.12 \pm 1.98$ | $-10.73 \pm 2.09$ |
| VIII | $-25.5$ | $-25.22 \pm 0.17$ | $-19.84 \pm 2.10$ | $-19.75 \pm 2.15$ |
| IX | $-17.4$ | $-14.47 \pm 0.53$ | $-12.21 \pm 1.65$ | $-13.40 \pm 1.87$ |
| X/SL+3 | $-23.2$ | $-20.94 \pm 0.32$ | $-19.19 \pm 2.92$ | $-19.99 \pm 2.15$ |
| XI/SL+4 | $-15.8$ | $-13.63 \pm 0.36$ | $-8.71 \pm 2.18$ | $-10.36 \pm 2.37$ |
| XII/SL+5 | $-23.2$ | $-21.74 \pm 0.34$ | $-19.43 \pm 2.44$ | $-20.42 \pm 1.49$ |
| XIII | $-17.7$ | $-16.13 \pm 0.18$ | $-12.74 \pm 2.11$ | $-13.86 \pm 1.72$ |
| XIV | $-31.0$ | $-29.36 \pm 0.40$ | $-21.80 \pm 2.42$ | $-22.09 \pm 2.76$ |

Table S4. Folding free energy of the 14 MS2 hairpin motifs together with the context of the native genome with length $L = 30$ nt. Listed are the minimum free energy of the motif sequence $F_{\text{motif}}$, the average free energy of the structural ensemble obtained for the motif sequence $\overline{F}_{\text{ensemble}}$, and the average free energies of the structural ensemble of the mono- and dinucleotide shuffled motif sequence ($\langle\overline{F}\rangle_{\text{mono}}$ and $\langle\overline{F}\rangle_{\text{di}}$, respectively). We produce 100 unique shuffles for each case where possible; due to the short motif sequence length, the number of dinucleotide shuffles is lower for motifs IX, XIII, and SL+5 (18, 12, and 84 dinucleotide shuffles, respectively).

[1] Ó. Rolfsson, S. Middleton, I. W. Manfield, S. J. White, B. Fan, R. Vaughan, N. A. Ranson, E. Dykeman, R. Twarock, J. Ford, *et al.*, J. Mol. Biol. **428**, 431 (2016).
[2] F. Pedregosa, G. Varoquaux, A. Gramfort, V. Michel, B. Thirion, O. Grisel, M. Blondel, P. Prettenhofer, R. Weiss, V. Dubourg, J. Vanderplas, A. Passos, D. Cournapeau, M. Brucher, M. Perrot, and E. Duchesnay, J. Mach. Learn. Res. **12**, 2825 (2011).
[3] G. J. McLachlan and S. Rathnayake, WIREs Data Min. Knowl. Discov. **4**, 341 (2014).
